# Warming treatments shift the temporal dynamics of diversity and composition of bacteria in wild blueberry soils

**DOI:** 10.1101/2024.10.03.616585

**Authors:** Oluwafemi A. Alaba, Suzanne L. Ishaq, Yu-Ying Chen, Lily Calderwood, Jianjun Hao, Yong-Jiang Zhang

**Author notes:** **Emails:** SI YYC LC JH. **Corresponding authors:** Oluwafemi A Alaba, Department of Botany and Plant Sciences, University of California, Riverside, Riverside, CA 92521, Yong-Jiang Zhang, School of Biology and Ecology, University of Maine, Orono, ME, 04469.

## Abstract

Soil bacterial communities are a crucial biological indicator of soil health and crop performance; however, their response to climate change remains poorly understood. In Maine, wild blueberry farms are experiencing unprecedented temperature changes, which may exacerbate microbial responses and potentially harm the crop. To elucidate the response of bacterial communities to warming during the growing season, we employed passive and active open-top chambers to simulate climate warming scenarios in wild blueberry fields. Warming treatments elevated atmospheric temperatures by 1.2 and 3.3 °C (passive and active warming), respectively, but did not affect soil temperatures. Nevertheless, soils in the active warming treatment exhibited significantly lower water content than ambient conditions. Overall, soil bacterial diversity and richness (June, July, and August data combined) under the warming (passive and active) treatments and ambient controls did not demonstrate significant differences after two years of experimental warming. However, significantly higher bacterial evenness and diversity under warming treatments were observed in the early growing season (June). Our study also reveals pronounced seasonal shifts in the evenness and diversity of bacteria in wild blueberry soil, suggesting that the variation in bacterial community structure may be more influenced by seasonal changes in temperature and plant activity during the growing season than by warming treatments. The increased bacterial evenness and diversity under warming treatments in June may be attributed to advanced plant phenology, indicating a potential future shift in seasonal dynamics of bacterial activity under global warming.

## 1. Introduction

Soil microorganisms play a crucial role in agricultural systems, serving as essential biological indicators of overall soil health, crop performance, and agricultural sustainability. These microbes establish symbiotic relationships with plant roots, obtaining sugars and organic acids from root exudates (Clark, 1949; Kumar et al., 2007). In addition to nutrient acquisition, soil microorganisms contribute significantly to ecosystem processes. They facilitate the depolymerization of organic compounds, recycle nutrients into plant-absorbable forms, enhance plant resilience to environmental stresses and even plant defense mechanisms (Burges, 1967; Yuan et al., 2018; Khan et al., 2019; Shah et al., 2021; Prasad et al., 2021). These functions underscore the fundamental role of soil microbes in supporting plant health and agricultural productivity. Agricultural management practices, including fertilization, mulching, tillage, and cropping systems, significantly influence soil microbial diversity and composition in both forest and agricultural environments (Ishaq, 2017; Madegwa & Uchida, 2021; Ishaq et al., 2020; Yeboah et al., 2016). Additional factors affecting soil microbial communities include soil pH (Zhalnina et al., 2015), vegetation type (Ravit et al., 2006; Hui et al., 2017), heavy metal contamination (Kandeler et al., 1996), land use (Kuramae et al., 2012), and soil water content (Brockett et al., 2012; Schimel, 2018). Recent research has increasingly focused on the detrimental effects of climate change on soil microbial communities and strategies for mitigating these impacts to improve soil health (Zogg et al., 1997; Balser et al., 2010; Mandal & Neenu, 2012; Ishaq et al., 2020).

Global temperature increases, averaging approximately 1.1°C, will rise by an additional 5.4°C by the end of the century if greenhouse gas emissions continue unabated (IPCC, 2014). These temperature changes are critical factors potentially affecting soil microbial communities, although a comprehensive understanding of their impact remains incomplete. Elevated temperatures can reduce soil moisture due to increased evapotranspiration, adversely affecting plant-microbe interactions (Rasmussen et al., 2020). For instance, long-term warming in temperate forests has already demonstrated significant alterations in soil bacterial community composition (DeAngelis et al., 2015). Furthermore, seasonal variation, which may result from a combination of environmental variables such as vapor pressure deficit, relative humidity, solar radiation, soil moisture, precipitation, and temperature, occasionally exerts a more pronounced effect on microbial community composition than temperature alone (Madegwa & Uchida, 2021; Pold et al., 2021). Different responses in microbial communities have been observed in short-term experiments based on initial soil properties and warming conditions (Zhao et al., 2022). Specific bacterial genera, such as *Pseudomonas* and *Bacillus*, can enhance plant yield and quality under water stress (Nordstedt & Jones, 2020; Paliwoda et al., 2022). *Acidothermus*, a thermophilic *Actinobacterium*, has also accelerated plant biomass decomposition in warmer environments (Viitamäki et al., 2022). However, elevated temperatures can also increase the prevalence of soil-borne pathogens, posing risks to plant health and agricultural productivity (Delgado-Baquerizo et al., 2020). Thus, understanding the responses and roles of soil microbes in maintaining the performance of perennial crops amidst rising temperatures remains a critical research challenge (Leisner, 2020).

Wild blueberries (mainly *Vaccinium angustifolium* Aiton), a perennial fruit crop native to North America, are grown on a 2-year cropping cycle, with berry harvests occurring every other year. Their rhizomes form a dense network of natural carpets that host diverse microorganisms within a thin layer above the sandy soil, with a pH of 4 (Li et al., 2020; Brown, 1966). The interactions between wild blueberries and their associated microorganisms are crucial for crop growth, yield, and fruit quality (Ramasamy et al., 2011; Li et al., 2020). The wild blueberry farm provides an ideal ecosystem for examining the effects of global warming on the microbial communities due to its distinctive soil structure and the prevalent climate warming impacting these growing fields (Schattman et al., 2020; Barai et al., 2021; Tasnim et al., 2021; Alaba et al., 2024). Many studies revealed that warming shifted soil microbial diversity and activity, suggesting that it could play roles in above-ground plant growth and yield (Zhang et al., 2005; Luo & Weng, 2011; Lladó et al., 2017; Chen et al., 2020). Climate warming alters the phenology, morphology, physiology, and fruit quality of wild blueberries (Chen, 2021; Alaba et al., 2024), and the economic viability of wild blueberry production in regions such as Maine and Canada is anticipated to face challenges due to the detrimental effects of warming (Yarborough, 1997; Drummond et al., 2009; Alaba et al., 2024). While previous studies have primarily focused on above-ground responses of wild blueberries to climate warming (Tasnim et al., 2021; Chen, 2021; Alaba et al., 2024), there is a pressing need to explore how below-ground microbial communities respond to current unprecedented climate warming.

This study aims to profile the bacterial community composition and diversity in wild blueberry fields under different climate warming scenarios and ambient conditions *in situ*, thereby providing the basis for potential soil management strategies that can sustainably lessen the impact of global warming in the future. Using Illumina amplicon 16S rRNA gene sequencing, we investigated the effect of warming treatments over two years (2019 and 2020) of growing seasons on soil bacterial communities. We hypothesized that elevated atmospheric temperatures of 1.2 °C and 3.3 °C would alter soil temperatures, reduce soil water content, and shift soil microbial community composition and abundance in open-top warming chambers compared to ambient control plots. Additionally, since the summer is warmer than the late spring of the growing season, we predicted that microbial community composition at the beginning of the growing season (June) would differ from that at the end of the growing season (July and August), regardless of warming conditions. Our findings aim to provide insights into the bacterial community responses to climate change and their implications for plant-microbe interactions and ecosystem functioning.

## 2. Materials and Methods

### 2.1. Study site and experimental design

The experiment was conducted at the Blueberry Hill Research Farm in Jonesboro, Maine (44.644° N, 67.646° W) in 2019 and 2020. The site has gravelly sandy loam soils with a pH of 4.7 and is characterized by a humid continental temperate climate (Smagula and Hepler, 1978; García-Gaines and Frankenstein, 2015). The average annual temperature at the site is 6.9 °C, and the average annual precipitation is 1298 mm (NOAA, 2023).

Open-top chambers (see the design in Tasnim et al., 2020) were installed in the field in April 2019, before the re-sprouting of wild blueberries from belowground biomass, to simulate realistic warming scenarios. Both active and passive heating open-top chambers were constructed with dimensions of 55 cm in height and 100 cm in width, inclined at 60 degrees on six sides to enclose wild blueberry plants using translucent polycarbonate sheets (Tasnim et al., 2020). The active heating (AH) chambers utilized heating tapes (Briskheat, Columbus, OH, USA) within the chambers to consistently elevate the mean air temperature by 3.3 °C for two years (2019 and 2020), while the passive (PH) chamber, lacking heating tapes, increased the mean air temperature by 1.2 °C. The ambient condition (CON) has no open-top chamber. A randomized block design with six genotypes was used. Specifically, six morphologically diverse wild blueberry (*V. angustifolium*) genotypes were randomly selected and employed as blocks containing the three distinct temperature treatments: AH, PH, and CON. Weather stations positioned approximately 10 cm above the soil were installed in the center of ambient plots and warming open-top chambers to monitor atmospheric conditions. Soil temperatures and soil volumetric water contents were determined using a TDR 150 Soil Moisture Meter (Spectrum Technologies Inc., Aurora, IL, USA) installed in the center of the plots at a depth of 5 cm for this study. To compare the effects of different heating treatments on soil temperature and water content, we performed a one-way ANOVA followed by a post-hoc Tukey’s HSD test to identify significant differences between treatments. No fertilizer or irrigation was applied during the two-year study period.

### 2.2. Soil sampling and processing

Soil sampling was conducted during the growing season of the second year of the experiment, 2020, on specific dates: June 09th (late spring), July 19th (early summer), and August 18th (late summer). A 3 cm diameter soil probe was used to collect three samples from the top 10 cm of soil at each treatment plot, removing the top of the organic matter. The fresh core samples were mixed thoroughly in the collection bags to create composite soil samples, which were then taken to the laboratory at the University of Maine. The composite soil samples were sieved using a 2-mm sieve to remove roots and stones before storing aliquots at -20 °C for DNA extraction.

### 2.3. Soil DNA isolation and real-time quantitative PCR (qPCR)

Soil and control samples were collected from three experimental treatments: active heating, passive heating, and ambient (control) conditions. Forty-eight soil samples and four control samples (one per extraction batch) were processed for DNA extraction. To ensure the selective amplification of DNA from viable cells, soil samples were diluted in sterile phosphate-buffered saline, homogenized, and treated with propidium monoazide (PMA; BioTium) at a final concentration of 25 μM, following the manufacturer’s instructions (Nocker et al., 2007). PMA binds covalently to DNA within dead cell membranes and free DNA, thereby preventing their amplification in downstream analyses. DNA was extracted from the PMA-treated soil samples (n = 48) and the no-template (water) controls (n = 4) using the Quick-DNA Fecal/Soil Kit (Zymo Research, Freiburg, Germany), which is designed explicitly for soil-based microbial communities. The quantity and purity of the extracted DNA were assessed using a Thermo Scientific™ NanoDrop™ OneC Microvolume UV-Vis Spectrophotometer (Thermo Scientific, Waltham, MA, USA). For microbial community analysis, the V3-V4 region of the 16S rRNA gene was amplified using the primers 515F and 926R according to the protocols established by The Earth Microbiome Project (Walters et al., 2016). The amplification products were sequenced on an Illumina MiSeq platform using the 2 x 300-nt V3 kit (Molecular Research Labs, Clearwater, TX, USA). Quantitative PCR (qPCR) was used to quantify the bacterial 16S rRNA gene copies in the soil samples. The qPCR analysis was performed using the Luna Universal qPCR Master Mix Kit (New England Biolabs, Ipswich, MA, USA) with fluorescence detection following the manufacturer’s protocol. Gene block standards from Integrated DNA Technologies (IDT) were serially diluted from 10^6 to 10^1 copies. The qPCR amplification utilized bacteria-specific primers 1048F Bac and 1194R Bac (Horve et al., 2020). The reaction mixture consisted of 2 μL of genomic DNA, 0.5 μL of each primer, 10 μL of SYBR Green qPCR master mix (New England Biolabs, Ipswich, MA, USA), and 7 μL of nuclease-free water, in a total volume of 20 μL. Amplification was conducted using an Applied Biosystems StepOne™ system (Applied Biosystems, Foster City, CA, USA) with the following thermal cycling conditions: an initial denaturation at 95 °C for 2 min, followed by 25 cycles of 95 °C for 30 s, 55 °C for 30 s, and 72 °C for 30 s, with a final extension at 72°C for 5 min. Each sample and standard dilution were analyzed in triplicate. The bacterial 16S rRNA gene copy number per μL of DNA was determined based on the linear relationship between the mean cycle threshold (Ct) values and the logarithm of gene copy numbers of the standards.

### 2.4. Amplicon sequencing analysis

Raw sequencing data from Illumina were processed using the DADA2 pipeline (version 1.16) within the R environment (v 4.1.3), following the procedures outlined by Callahan et al. (2016). The DADA2 pipeline applied standard parameters for several critical steps: trimming sequences for quality control (trimLeft = 15, trimRight = 20, truncQ = 2, maxEE = 2, maxN = 0, rm.phix = TRUE), denoising data, dereplicating reads, merging paired-end reads, and filtering out chimeric sequences using the de novo algorithm built in. Taxonomic identification of the resulting amplicon sequence variants (ASVs) was achieved using the SILVA reference database (version 138). To ensure data integrity, ASVs from negative control samples, which underwent the same DNA extraction and sequencing procedures as the soil samples, were used to remove potential contaminants from the soil data. Additionally, sequences classified as chloroplasts and mitochondria were excluded from the analysis. A rarefaction curve was generated and visualized using R to assess the sequencing depth. To standardize the data and prevent biases in clustering analyses and diversity metrics, samples were scaled and evened with the lowest sequencing depth of 13,300 reads. The final ASV table and corresponding taxonomy file were utilized for subsequent microbial diversity and statistical analyses.

### 2.5. Microbial diversity

The rarefied phyloseq object, as detailed by McMurdie and Holmes (2013), was utilized to estimate Shannon indices, which measure alpha diversity, encompassing bacterial ASV richness and evenness. The Mann-Whitney-Wilcoxon non-parametric test assessed the normality of ASV abundance and distribution across different treatments (p < 0.05). Visualizations were created using boxplots and correlation matrices to illustrate the top 20 most abundant ASVs, with plots generated through the ‘ggplot2’ and ‘corrplot’ functions. Beta diversity was analyzed using Bray-Curtis dissimilarity metrics, and constrained correspondence analysis (CCA) was performed to evaluate the effects of warming treatments and seasonal changes on bacterial community composition. Permutational multivariate analysis of variance (PERMANOVA) was conducted to determine significant differences in bacterial composition between warming and seasonal change groups. A two-way ANOVA was used to analyze the effects of warming treatments and seasonal variations on soil bacterial diversity and composition, followed by post-hoc comparisons using Fisher’s Least Significant Difference (LSD) test (p ≤ 0.05). All statistical analyses were executed in R using RStudio version 2022.07.1+554 (R Team; Allaire, 2012). The raw sequence data have been deposited in the NCBI Sequence Read Archive (SRA) under Bioproject accession number PRJNA925843.

## 3. Results

### 3.1. Effect of warming on soil temperature and water content

The experimental warming using open-top chambers elevated atmospheric temperatures by 3.3°C in the active warming chambers and by 1.2°C in the passive warming chambers compared to ambient control plots during the wild blueberry growing seasons (Chen et al., 2022). Despite these atmospheric temperature increases, the warming treatments did not significantly change average soil temperatures, as no differences were observed among the treatment groups (Fig. 1a). However, the soil volumetric water content exhibited significant variations across treatments. In August, soil moisture was markedly lower (p < 0.001) in the active warming chambers compared to both the passive warming and ambient control plots (Fig. 1b). Additionally, during May and June, soil volumetric water content was significantly reduced (p < 0.05) in the active warming chambers relative to the ambient control plots (Fig. 1b).

**Fig 1.**
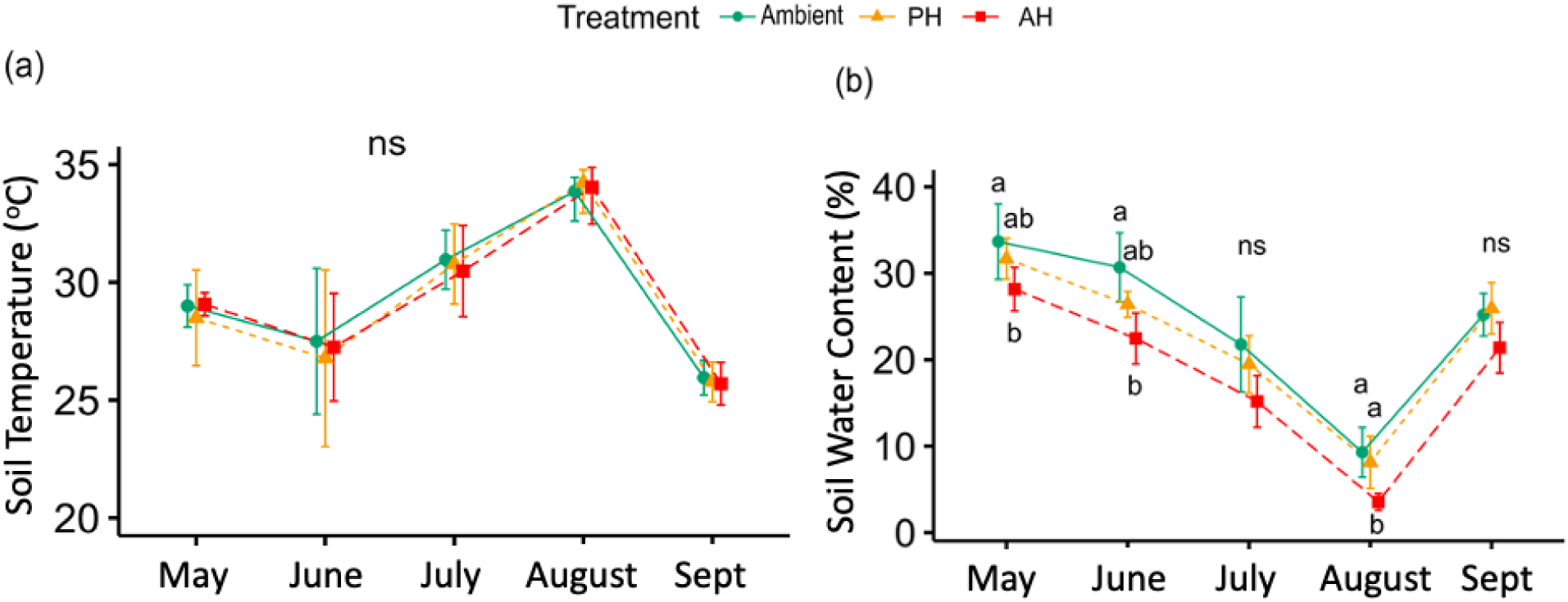
Comparison of average soil temperature (a) and volumetric water content (b) in wild blueberry soils under ambient control (green dots), passive warming (PH) treatments (orange triangles), and active warming (AH) treatments (red squares). “ns” denotes no significant differences among treatments (p > 0.05). Different letters on the dots indicate significant differences between treatments determined by Tukey’s HSD post hoc test following analysis of variance (p < 0.01 in August and p < 0.05 in May and June).

### 3.2. Soil bacterial richness, evenness, and diversity under warming conditions

A total of 28,877,870 raw 16S rRNA reads were obtained from 52 soil samples exposed to simulated warming treatments and ambient conditions. After quality control, an average of 301,501 high-quality sequence reads were retained. From the initial 113,120 bacterial sequence variants (SVs), 38,432 chimeric sequences were identified and removed, resulting in a refined dataset of non-chimeric and rarefied amplicon sequence variants (ASVs). Subsequent analysis of overall bacterial richness, Shannon evenness, and diversity revealed no significant differences between warming treatments (active and passive) and ambient conditions when data from different months were combined (Fig. 2a & b). However, bacterial richness and diversity significantly increased with seasonal changes throughout the growing season (Fig. 2c & d). The observed richness and diversity peaked in August and exhibited a statistically significant difference (p ≤ 0.0001) compared with June under combined treatements.

**Fig 2.**
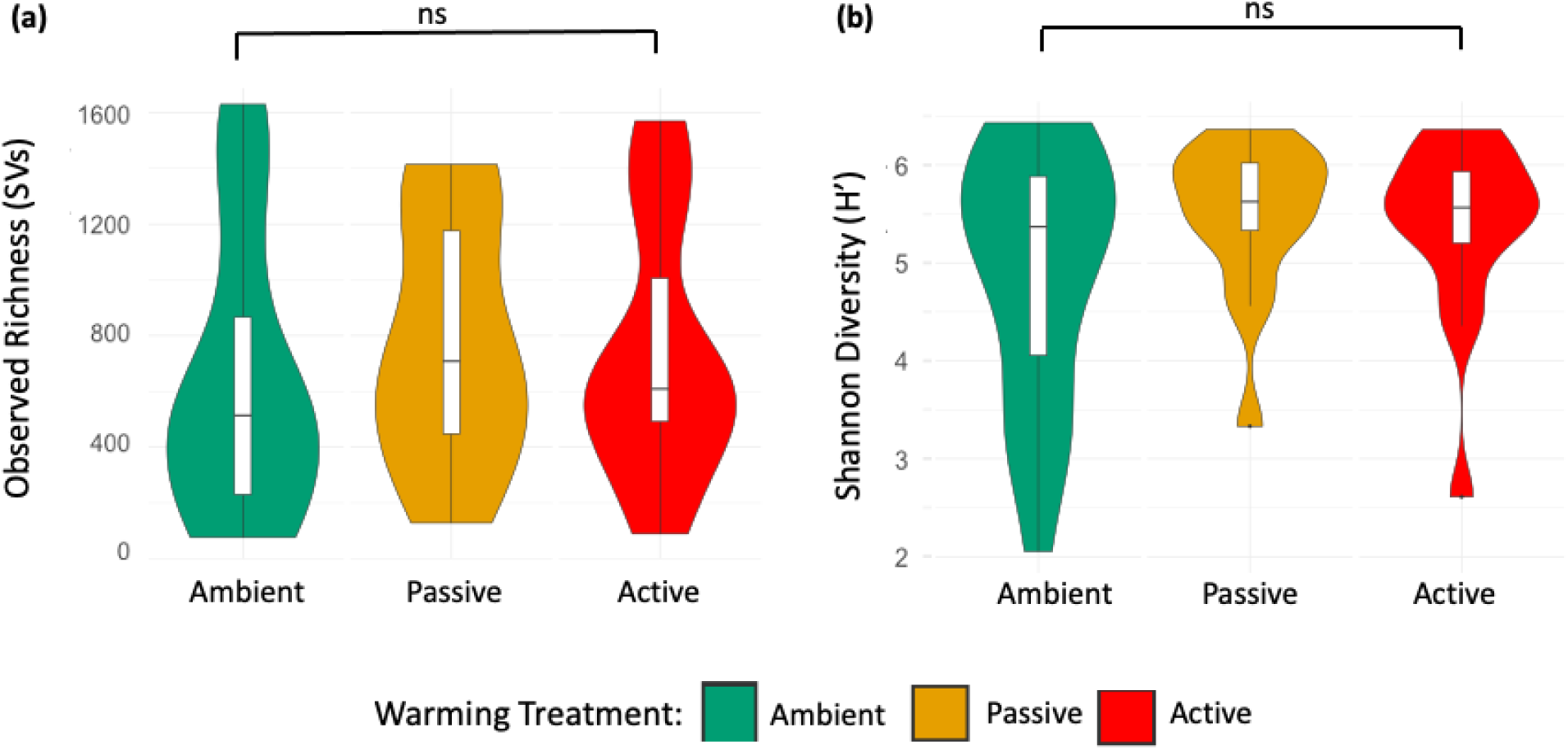
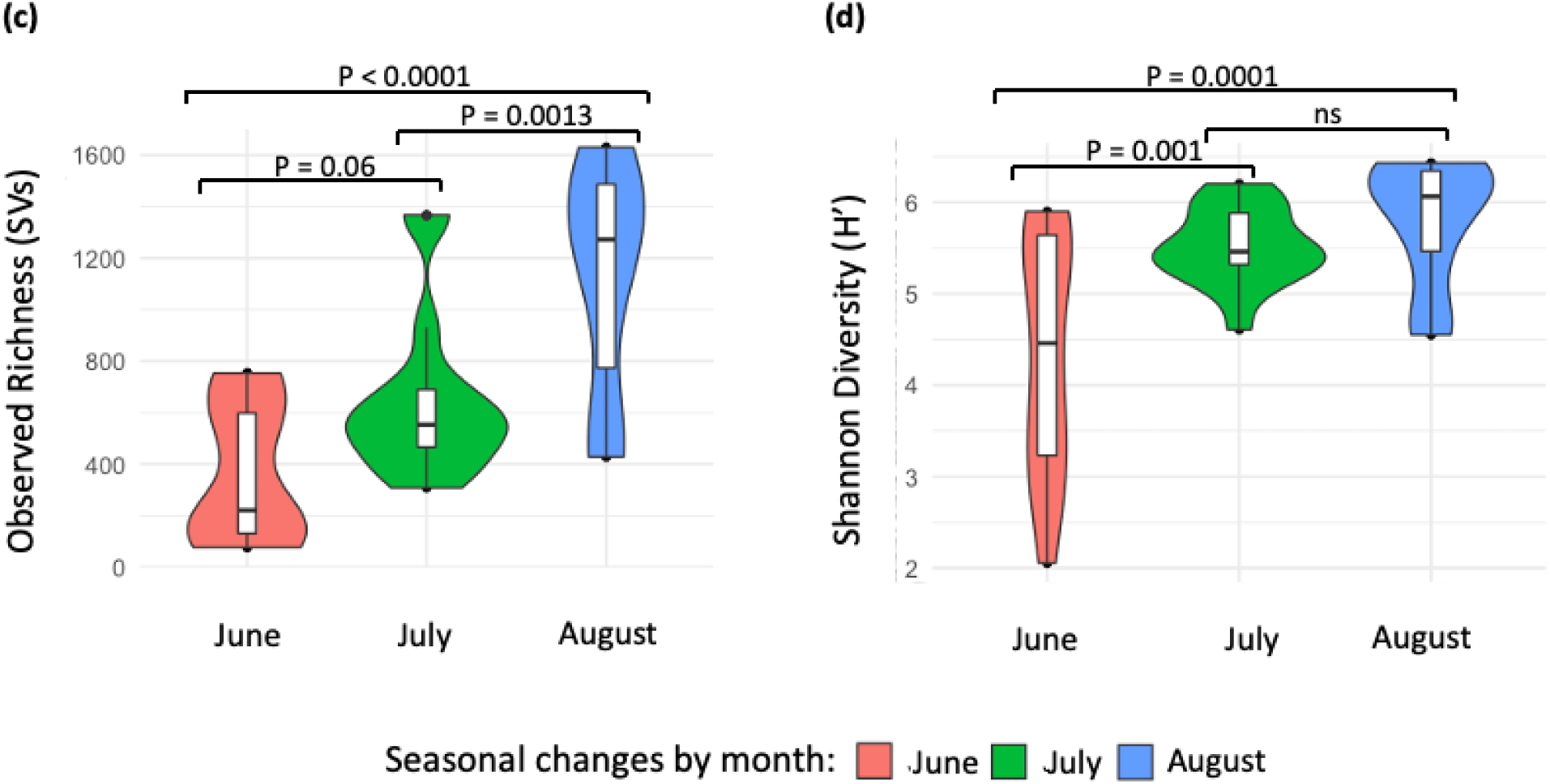
Comparison of overall bacterial community richness (a) and Shannon diversity (b) in wild blueberry soils under warming treatments (passive and active) and ambient control. There were no significant differences between warming treatments and the ambient control (p > 0.05). Panels (c) and (d) show observed richness and Shannon diversity of soil bacteria by season—late spring (June 9), early summer (July 19), and late summer (August 18) – under combined treatments. Significant differences were observed between spring and summer under combined conditions (p < 0.05).

The analysis of the data revealed a notable seasonal difference in the overall richness, evenness, and diversity (Shannon-Weiner diversity index) of the bacterial community in the wild blueberry soil (Fig. 3). Specifically, we observed that bacterial richness and diversity were significantly higher in August compared to June, while no significant differences were found among the treatments in July and August (summer months). However, within the spring month (June), higher bacterial evenness and Shannon diversity were found in both the active and passive heating treatments compared to control (ambient) conditions (Fig. 3b & c). These findings indicate that the observed changes within the spring month are primarily attributed to the effect of warming conditions.

**Fig 3.**
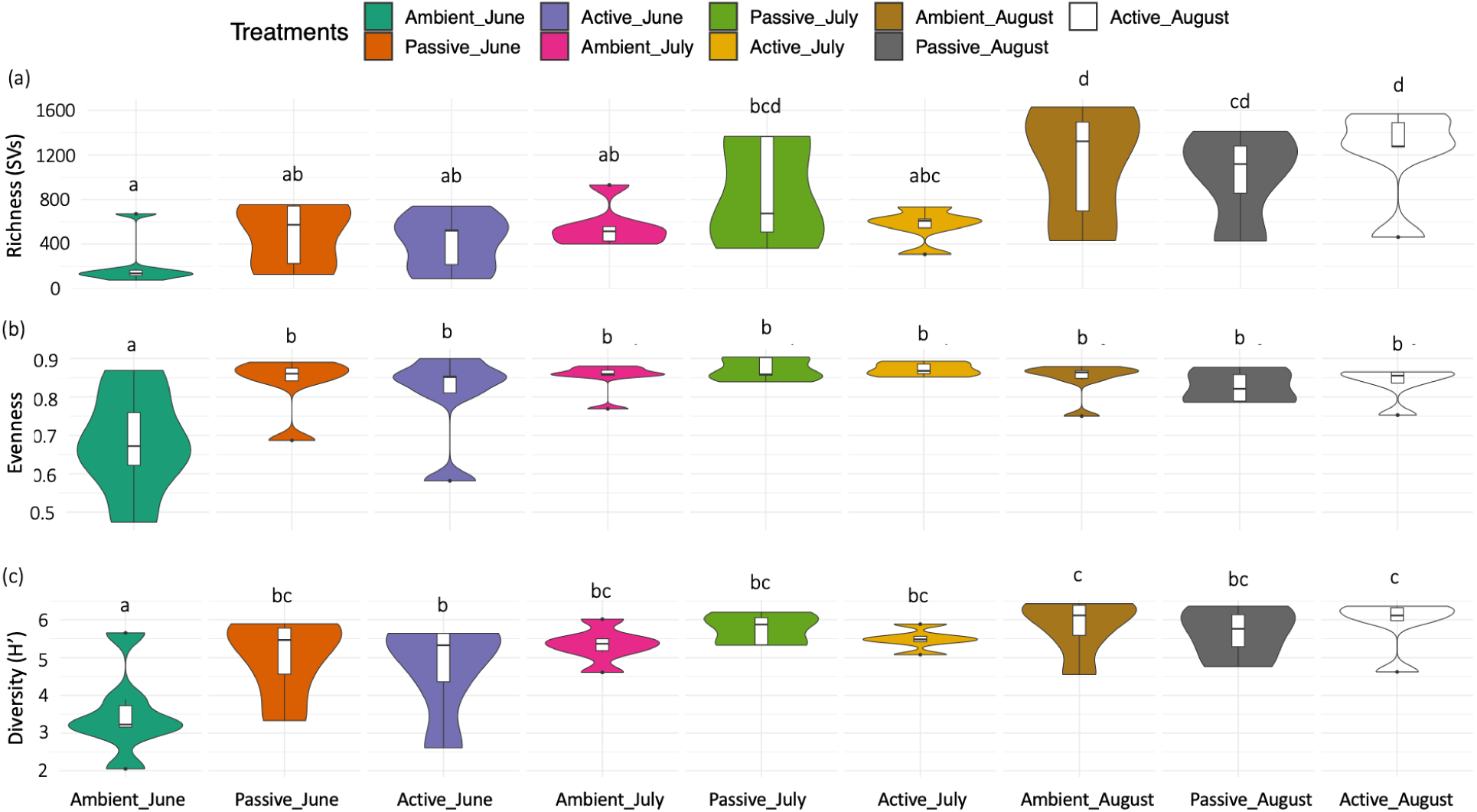
Comparison of bacterial community metrics in wild blueberry soil under climate warming treatments (Passive and Active) versus ambient control across spring and summer. (a) Observed richness (SVs), (b) Evenness, and (c) Diversity (Shannon index). The thick bar within each violin plot represents the mean. Letters indicate significant differences as determined by Tukey’s HSD post hoc test following analysis of variance, with significance tested at an alpha level of 0.05.

### 3.3. qPCR analysis of bacterial gene copy numbers under warming conditions

We quantified 16S rRNA gene copy numbers using qPCR to assess bacterial abundance in soil samples under ambient and warming conditions (Fig. S1). The results revealed no significant differences in bacterial populations, as indicated by consistent log10-transformed 16S rRNA gene copy numbers across both conditions. Interestingly, this pattern mirrors our amplicon sequencing results, which showed no significant changes in bacterial observed richness or Shannon diversity between warming treatments (passive and active) and ambient control soils (Fig. 2a & b). This consistency is likely due to the exclusion of seasonal variations from the qPCR analysis.

### 3.4. Bacterial relative abundance and composition under warming conditions

At the phylum level, the relative abundance of dominant bacterial phyla, as determined by 16S rRNA gene sequencing, exhibited variation across seasons and warming treatments (Fig. 4). The taxonomic analysis revealed that seasonal transitions from spring to summer had a significant impact on the composition and relative abundance of various bacterial phyla. The *Proteobacteria* phylum remained consistently abundant across all temperature treatments and seasons. Similarly, the *Verrucomicrobiota*, *Acidobacteriota*, *Actinobacteriota*, and *Planctomycetota* phyla exhibited high relative abundance across most treatments. However, in the Ambient June treatment, a marked reduction or near absence of these phyla was observed, indicating a distinct bacterial community structure (Fig. 4). *Firmicutes* and *Bacteroidota* were more abundant in spring compared to summer, while *Chloroflexi* exhibited a higher relative abundance in the summer samples. Finally, the abundance of *Bacteroidota* in spring was significantly lower under warming conditions than in the ambient control (Fig. 4).

**Fig 4.**
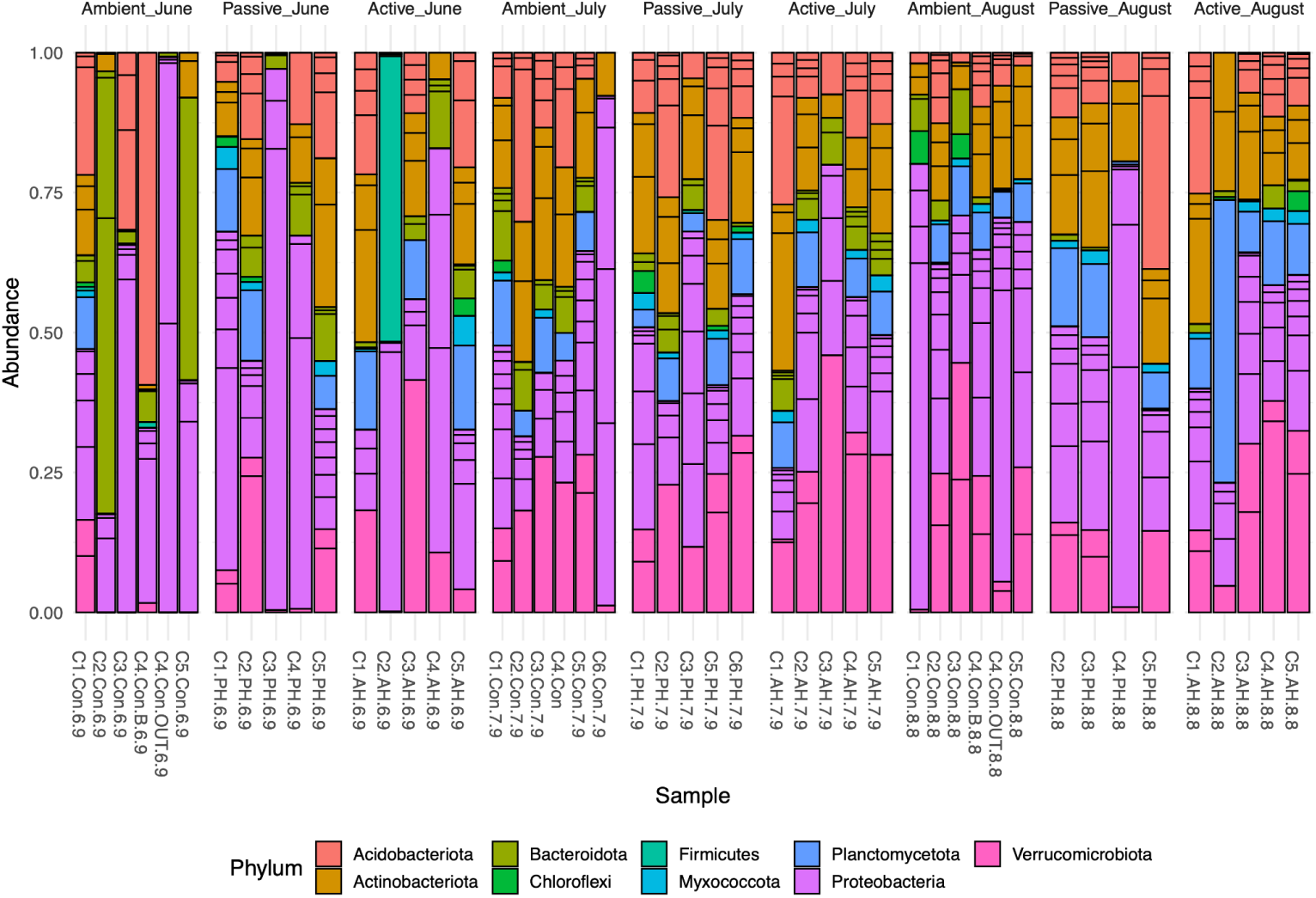
Relative abundance of bacterial phyla in wild blueberry soils under different treatments (ambient, active warming, passive warming) across the growing season (June to August).

Community structure is maintained by abundant bacteria. This study identified twelve predominant genera as having the highest relative abundance of amplicon sequence variants (ASVs) at finer resolution (Fig. 5). *Acidibacter* was the most abundant and displayed significant variability across seasons and warming treatments. *Sediminibacterium* was more prevalent in the control treatment than the warming treatments during spring. Other abundant genera included *Aquisphaera*, Candidatus *Xiphinematobacter*, Ellin6067 (an unclassified Betaproteobacteria), and *Roseiarcus* (Fig. 5).

**Fig 5.**
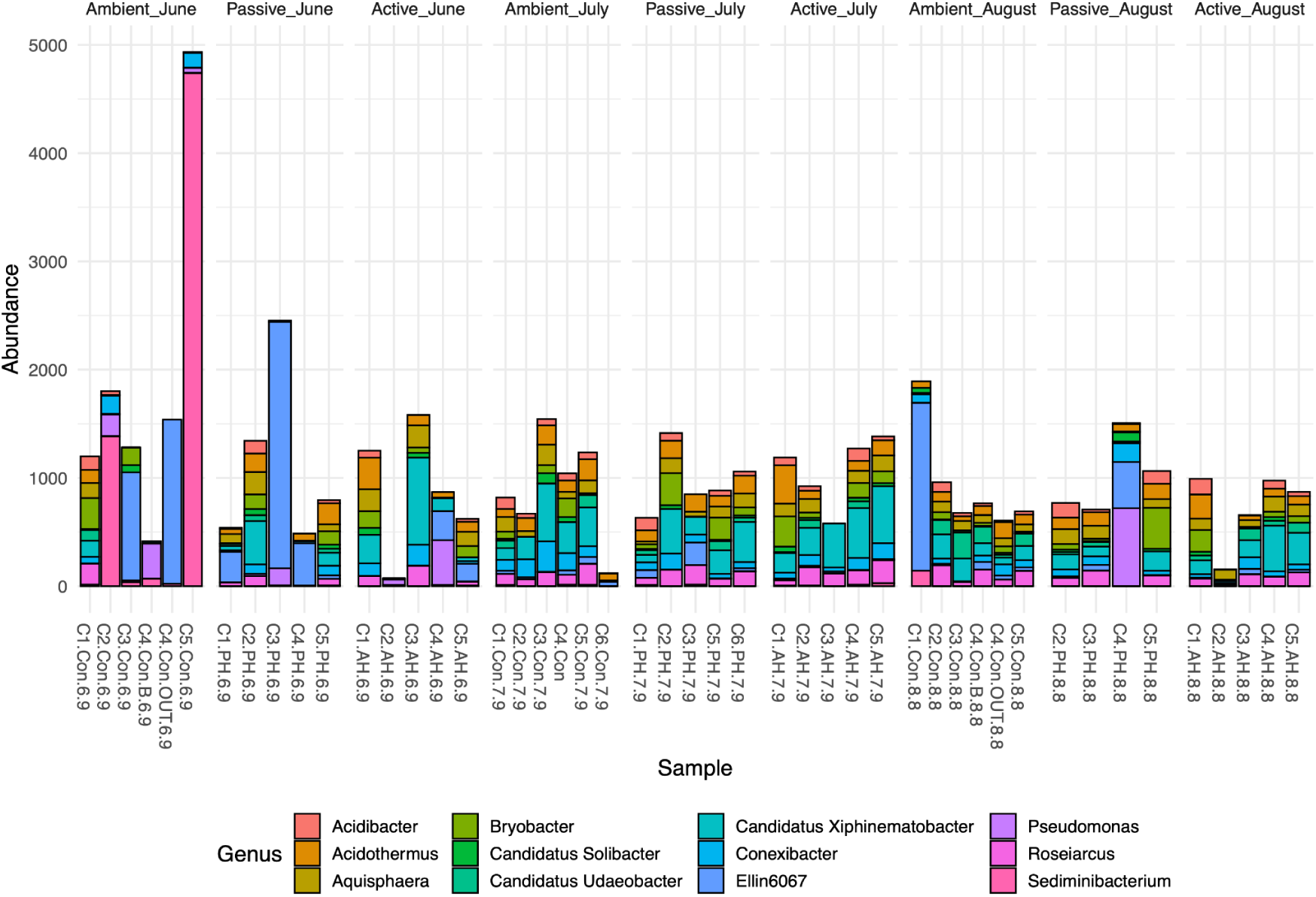
Abundance of the top 12 bacterial ASVs at the genus level in wild blueberry soils under different treatments (ambient, active warming, passive warming) across the growing season (June to August).)

Seasonal changes significantly influenced the taxonomic shifts in genus composition observed in our study. We further examined the relative abundance of twelve predominant genera—*Acidothermus*, *Aquisphaera*, *Bryobacter*, Candidatus *Xiphinematobacter*, *Conexibacter*, the candidate *Ellin6067*, *Flavobacterium*, *Methylocapsa*, *Roseiarcus*, *Sediminibacterium*, *Sphingobium*, and *Variovorax*—to assess variations between warming treatments and the ambient control. In June, the mean relative abundances of *Acidothermus*, *Aquisphaera*, and *Roseiarcus* were lowest in the ambient temperatures but increased consistently under warming conditions (Fig. 6). Conversely, the mean abundance of Candidatus *Xiphinematobacter* rose from June to July, peaking under the Active treatment in July before decreasing in August. *Ellin6067*, *Sediminibacterium*, and *Variovorax* were more abundant in the ambient control in June than in other treatments. *Flavobacterium*, *Methylocapsa*, and *Sphingobium* exhibited some outliers but no notable differences in abundance under the ambient control in June.

**Fig 6.**
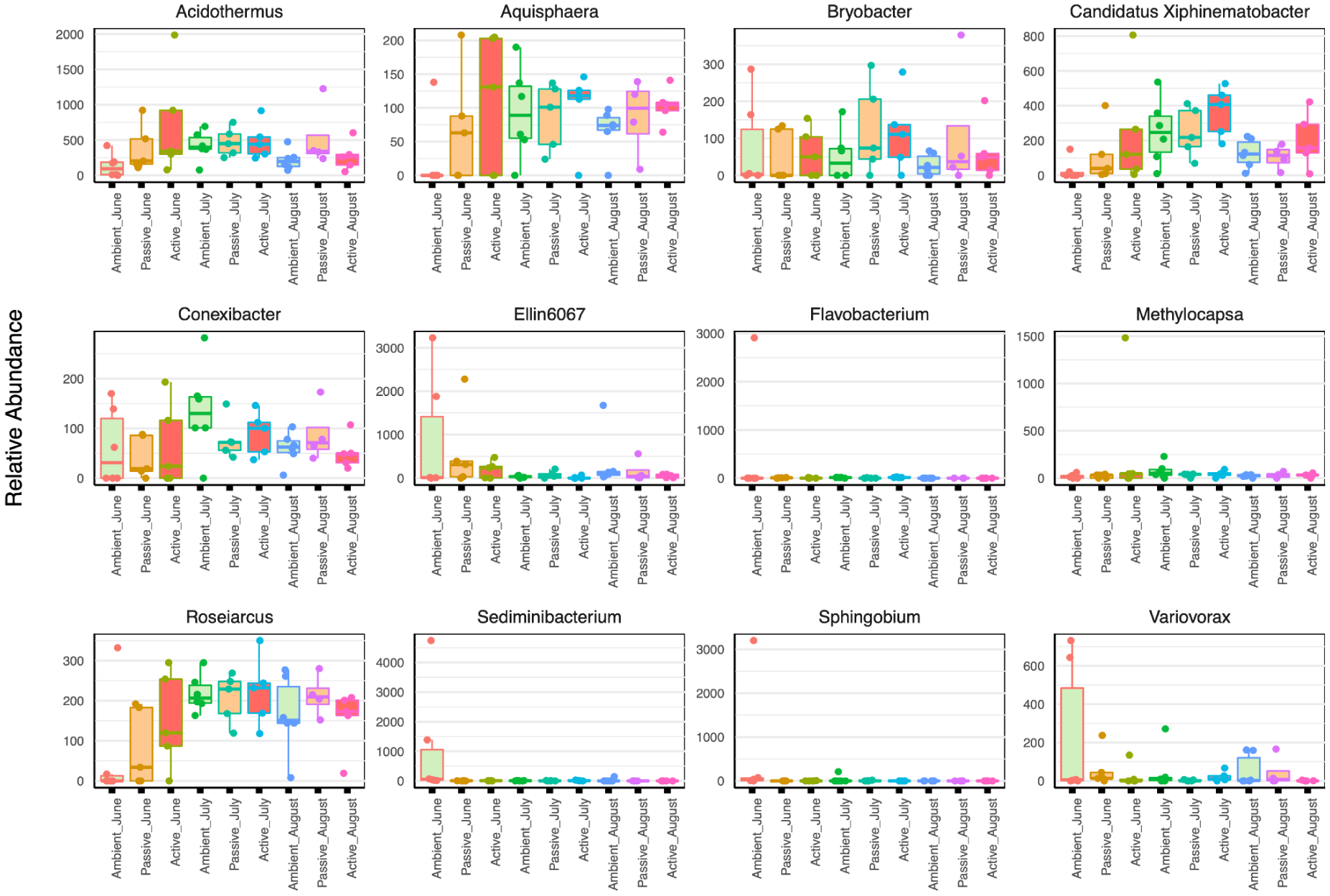
Boxplots illustrate the differences in the relative abundance of wild blueberry soil bacteria at the genus level under warming treatments and across seasonal changes from spring to summer. Data are presented for June (Ambient, Passive, Active), July (Ambient, Passive, Active), and August (Ambient, Passive, Active).

### 3.5. Predictor bacterial taxa under elevated temperatures and seasonal changes

Using a random forest model, we identified the top 30 discriminant bacterial taxa based on sequence variant estimates with significant importance (p < 0.05) across different seasons under warming treatments (Fig. 7). This analysis highlighted which bacterial taxa were most predictive of elevated temperatures in wild blueberry soil as the seasons transitioned from spring to summer. Key SVs in genera such as *Variovorax*, *Acidothermus*, *Bryobacter*, Candidatus *Xiphinematobacter*, and *Aquisphaera* were crucial predictors of bacterial abundance under warming conditions. Other significant predictors included *Gemmataceae*, *Flavobacterium*, and *Acidibacter*. The relative abundance of *Gemmataceae* and *Acidibacter* was high in summer but very low in spring, while *Flavobacterium* exhibited the opposite trend (Fig. 7). The model’s out-of-bag error rate was 76.6%, indicating a high level of uncertainty in differentiating between treatment groups. Nonetheless, the estimated error rates for ambient plots in June and July were 16% and 33%, respectively, suggesting that bacterial communities in these plots were more distinct from other treatment groups and important taxa could be identified with higher confidence scores.

**Fig 7.**
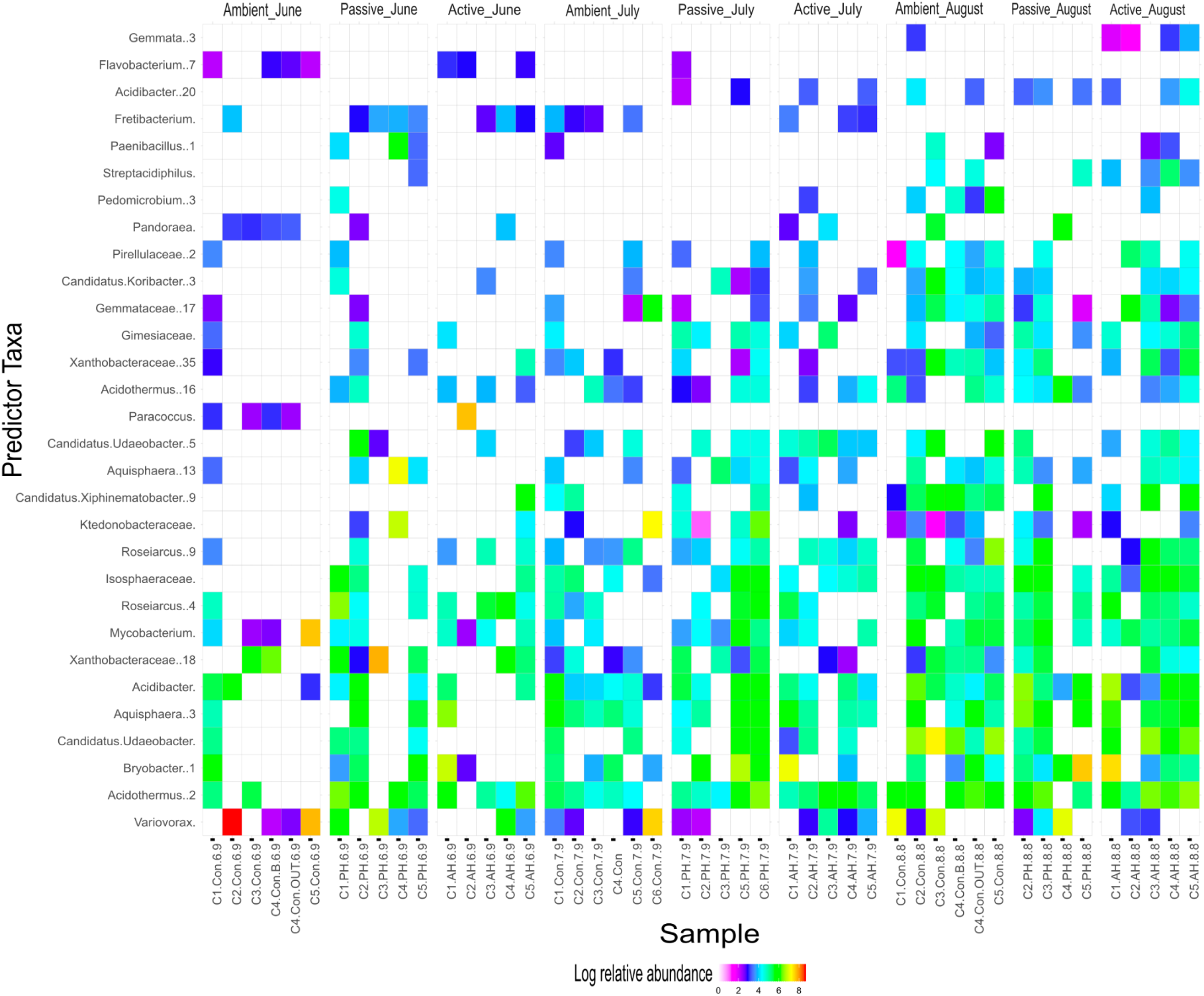
Heatmap generated using a permutational random forest model, showing the log relative abundance of the top 30 predictor taxa in wild blueberry soils under different temperature treatments (ambient, active, and passive) during climate warming in spring and summer. White blocks indicate where discriminant taxa were absent or present at very low levels. The model’s out-of-bag error rate was estimated at 76.6%.

### 3.6. Effects of seasonal changes and warming on bacterial community composition

We performed a Canonical Correspondence Analysis (CCA) using the Bray-Curtis dissimilarity method to explore the structure of soil bacterial communities during the growing seasons (spring and summer) under three temperature treatments. The CCA results indicated that the first and second axes explained 4.3% and 2.6% of the total variation in bacterial community structure, respectively (Fig. 8).

**Fig 8.**
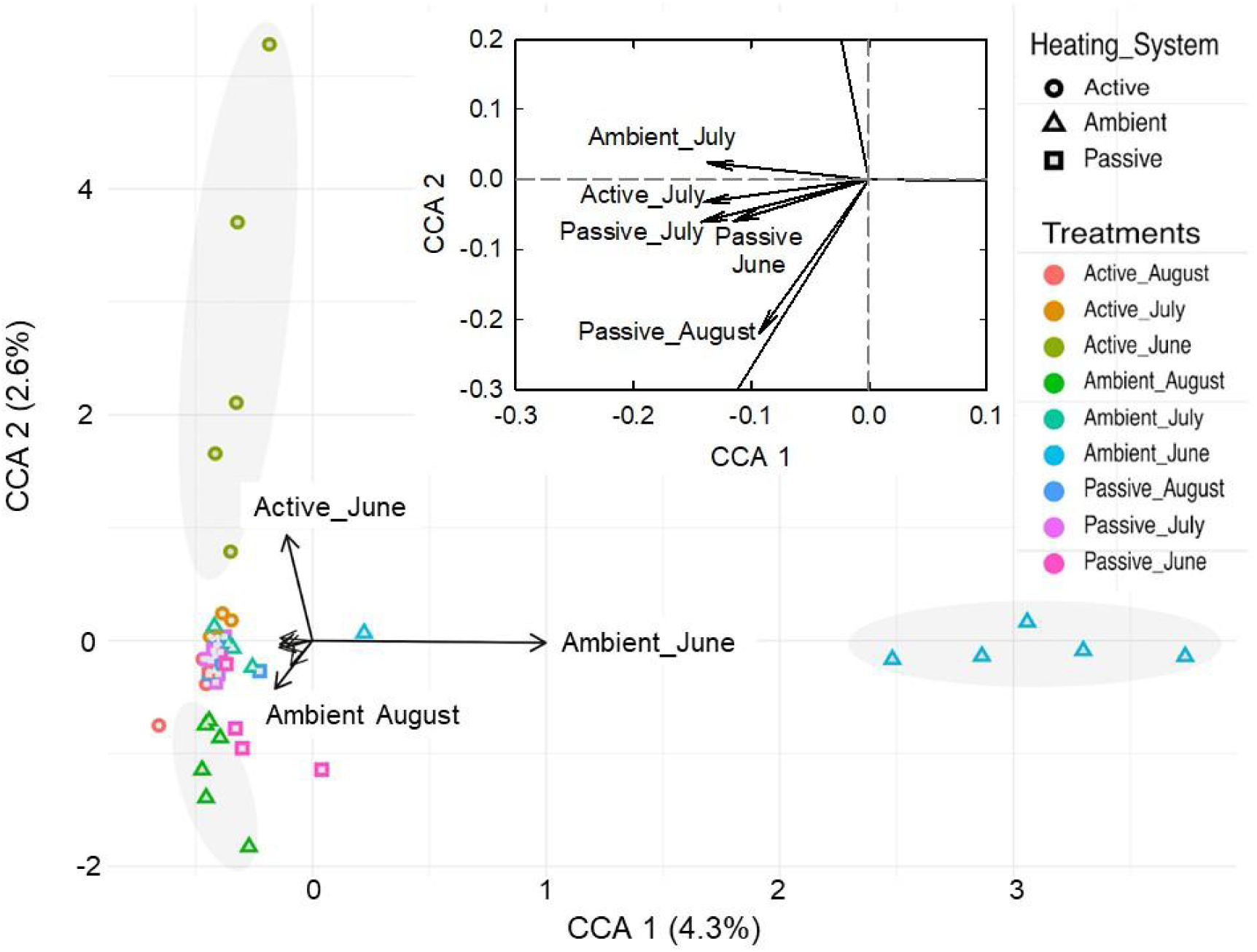
Constrained Correspondence Analysis (CCA) displaying the dissimilarity in bacterial community composition in wild blueberry soils based on the Bray-Curtis method, comparing climate warming treatments with ambient controls across different months of the growing season. PERMANOVA was used to assess significant differences between treatments. The inset provides an enlarged view of the central region of the figure.

The first axis of the CCA separated bacterial communities from ambient conditions in June from those in August, highlighting significant clustering of amplicon sequence variants (ASVs) primarily due to seasonal changes from June to August (Fig. 8). This separation suggests that the transition between these seasons strongly influences bacterial community composition. Moreover, the projected lengths and directions of the vectors demonstrated a positive relationship between bacterial community composition and the temperature treatments, revealing that bacterial communities in wild blueberry soils during the spring season (June) differed significantly between ambient control and warming treatments (both passive and active). These findings indicate that warming treatments have a more pronounced effect on soil bacterial communities during the early growing season.

Bacterial communities under warming treatment in July clustered closely with ambient controls in July, evidenced by Bray-Curtis dissimilarities, indicating that the passive treatments are weak to cause any substantial changes in the structure of the bacterial communities (Fig. 8). Interestingly, the bacterial communities in passive June showed a notable similarity to those in passive July and August. All these observations suggest that soil bacteria communities may exhibit both sensitivity and adaptability to warming treatments across different seasons.

### 3.7. Potential relationship between abundant bacteria taxa and diversity metrics

A Spearman’s rank correlation was performed to examine the relationships between the relative abundances of bacterial genera and alpha diversity parameters in wild blueberry soil. This analysis provides insight into the strength and direction of associations between different bacterial taxa (Fig. 9).

**Fig 9.**
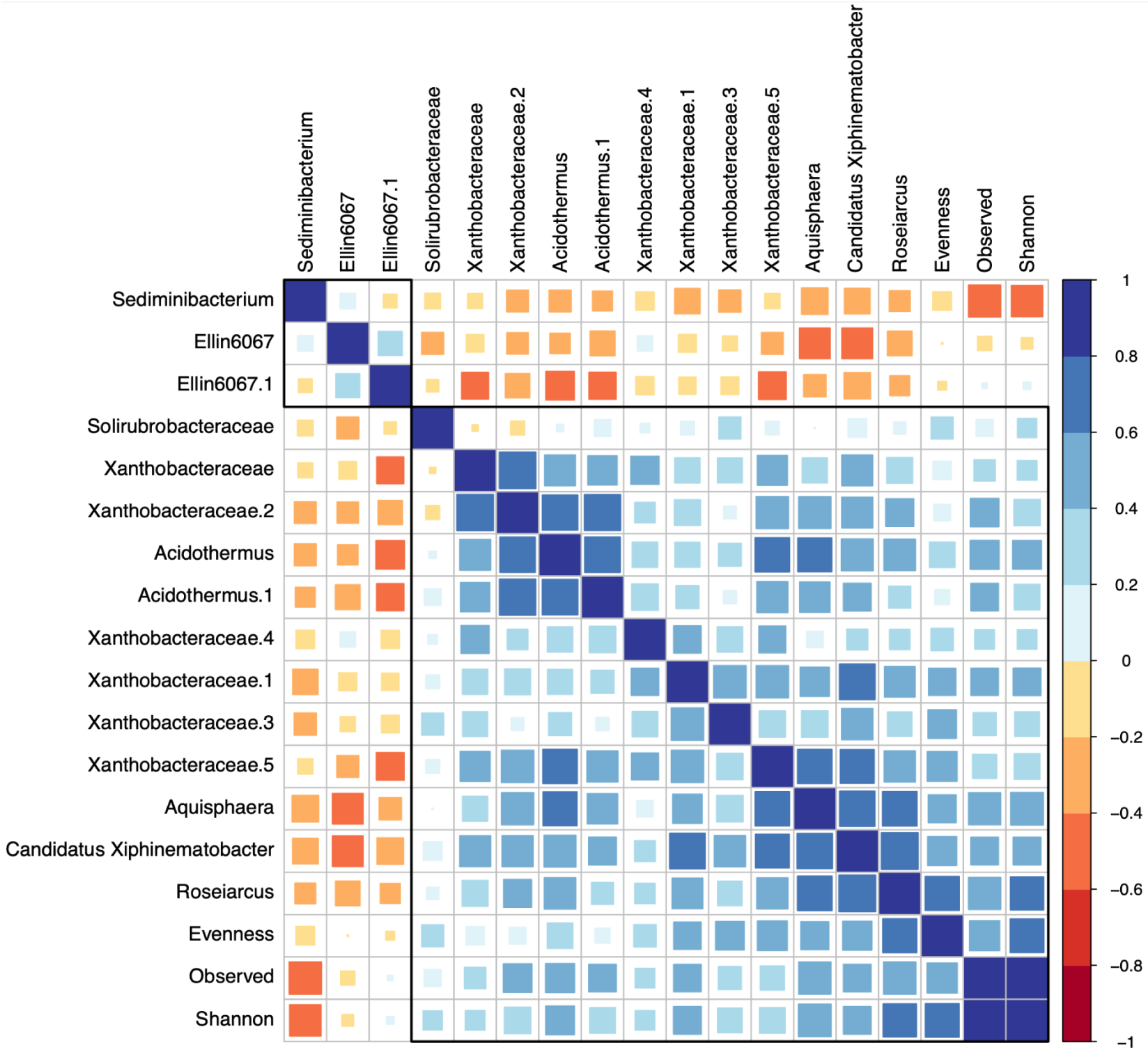
Spearman’s rank correlations between alpha diversity metrics and amplicon sequence variants (ASVs) of major bacterial taxa. White tiles indicate no correlation, light blue to blue tiles represent positive correlations, and orange to red tiles represent negative correlations. P-values were adjusted to < 0.05.

The study found a positive correlation between an SV in the genus *Acidothermus* and SVs in the family *Xanthobacteraceae*, as well as genera *Aquisphaera*, Candidatus *Xiphinematobacter*, and *Roseiarcus*, suggesting cooperative interactions among these taxa that may enhance the bacterial community structure in the soil. In contrast, the genus *Sediminibacterium*, prevalent in ambient temperature conditions during the spring, exhibited a negative correlation with alpha diversity parameters (Shannon diversity, observed richness, and evenness) and other abundant taxa, such as the candidate Ellin6067, an SV in the families *Solirubrobacteraceae* and *Xanthobacteraceae*, and SVs in the genera *Acidothermus*, *Aquisphaera*, Candidatus *Xiphinematobacter*, and *Roseiarcus*. This negative correlation suggests a competitive relationship between *Sediminibacterium* and other bacterial taxa (Fig. 9).

## 4. Discussion

Wild blueberry is a critical small fruit crop; however, limited knowledge exists regarding the soil bacterial communities in its non-amended soils, particularly under the influence of climate change. This study represents the first investigation of these bacterial communities during the growing season utilizing 16S rRNA amplicon sequencing and qPCR. Over a three-month period, we evaluated the effects of short-term *in situ* warming (1.2 to 3.3°C) on bacterial diversity, richness, and community structure. Despite increased soil water deficits due to warming, no significant alterations in bacterial diversity or richness were observed. Our hypothesis that warming would substantially alter the overall bacterial richness and diversity was unsupported, presumably due to the short duration of warming and the magnitude of heating that did not significantly raise the soil temperature in warming plots as anticipated. This same lack of effect over the short-term has been observed in other studies using open-top chambers and other crop species (Ishaq et al. 2020; Ouverson et al. 2022). However, we did find increased bacterial evenness and diversity under warming treatments in the early growing season, which may be attributed to advanced plant phenology. This suggests a potential shift in the temporal dynamics of bacterial activity in the future.

Our study, which observed no significant changes in bacterial community structure under continuous two-year warming, contradicts a recent study by Weedon et al. (2023) that reported substantial shifts in bacterial composition after four years of warming. Their experiment raised plot temperatures to 40°C, increasing soil temperatures by approximately 9°C, which led to a significant decrease in alpha diversity in the warming plots compared with controls (Weedon et al., 2023). Furthermore, numerous soil bacterial species formed the core taxa that can adapt effectively to changing environments due to their metabolic flexibility and physiological resilience (Meyer et al. 2004; Allison and Martiny 2008). This adaptability enables bacterial communities to resist changes despite short-term environmental stresses.

### 4.1. Soil bacterial diversity increased under warming in the early growing season

In June, bacterial evenness and diversity increased significantly under warming conditions, indicating that warming can influence microbial communities during the cooler season. This observation aligns with previous studies conducted in cooler regions, where warming has been demonstrated to enhance bacterial diversity (Fierer et al., 2003; Ishaq et al., 2020; Sheik et al., 2011). However, this warming effect during the warmer months of July and August was limited as no significant differences in evenness and diversity were observed between warmed and control plots. This suggests that while warming may enhance bacterial diversity and activity during temperature-limited seasons, its effects are moderated in the late growing season, where naturally elevated summer temperatures contribute to more stable bacterial communities. In line with our findings, previous studies demonstrated that seasonality might obscure the effects of warming, as bacterial communities were more strongly influenced during the early growing season compared to the mid-and late-growing seasons (Hu et al. 2023; Ishaq et al. 2020; Slaughter et al. 2015).

### 4.2. Bacterial composition and structure in response to warming and seasonality

The structure and composition of bacterial communities are crucial in performing roles such as nutrient cycling and decomposing organic matter in soils (Burges, 1967; Yuan et al., 2018). However, bacterial structure and function are influenced by environmental factors such as soil pH, moisture, and vegetation (Zhalnina et al., 2015; Hui et al., 2017). Our study observed shifts in community structure at the phyla level, particularly during the early growing season, when the warming effect is more pronounced in the cooler early season (June) compared with the warmer late season (August).

The bacterial phyla more abundant in warming plots in June include *Verrucomicrobiota*, *Planctomyoetota*, and *Firmicutes*, while *Bacteroidota* is more abundant with more than 50% in two of the samples in the Ambient June treatment. *Verrucomicrobiota*, which are involved in organic matter metabolism and soil fertility enhancement, exhibited fluctuations in relative abundance in response to environmental factors such as temperature (Islam et al., 2008). The increased abundance of *Verrucomicrobiota* during the summer months in our study may be attributed to the seasonal variation in ambient temperature. Thermoacidophilic strains of *Verrucomicrobiota* have been observed to be abundant under warmer conditions (Islam et al. 2008).

*Acidobacteriota*, *Actinobacteriota*, and *Proteobacteria* were consistent with those previously reported in wild blueberry soils (Yurgel et al., 2018). *Acidobacteriota* was abundant across all conditions, reflecting its resilience and essential role in regulating biogeochemical cycles (Kalam et al., 2020). *Actinobacteriota* phylum has been described as displaying adaptive strategies to environmental stresses, including warming-induced drought (Sheik et al., 2011; Hu et al., 2023). Therefore, we assumed that the insignificant effect of warming on the bacterial community in summer, as observed in this study, may result from bacterial resilience to small perturbations or disturbances, as short-term warming effects may be limited in their ability to shift bacterial communities. *Actinobacteriota*, known for its heat tolerance, was prevalent in July and August, underscoring its capacity to adapt to short-term warming conditions (Sheik et al., 2011). Overall, our findings emphasize the importance of accounting for seasonal dynamics when evaluating the effects of warming on microbial communities. While short-term warming may not induce large-scale changes in bacterial diversity, it can influence community composition and the functional roles of key phyla, particularly during cooler seasons.

Divergence in the beta diversity of bacteria communities corresponds to the response to changes in bacterial richness and alpha diversity (Rui et al. 2015). According to previous studies, warming affects bacterial richness, and alpha diversity changes the growing season more than in the mid-and late seasons, which is consistent with our results (Slaughter et al., 2015; Hu et al., 2023). The composition of bacterial communities in wild blueberry soils in June (spring, early growing season) and August (summer, late growing season) were different under ambient conditions, revealing the substantial effect of seasonal variations in microbial composition (Fig. 8). Under the warming treatments, the bacterial community composition was not similar to that of the ambient control treatment (Fig. 8). The effect of warming on bacterial communities in the early growing season may be more significant than in the late seasons, according to our study.

### 4.3. Key bacteria taxa and their association with diversity index

The abundances of specific bacterial taxa may be considered biological indicators in the transition from ambient to warming conditions across seasons. Certain bacterial genera, such as *Acidothermus* and Candidatus *Xiphinematobacter*, exhibited higher abundance in warmed soils, particularly in spring. The increased presence of *Acidothermus*, a cellulolytic bacterium, suggests that warming may enhance plant biomass degradation, thereby enriching the soil with bioavailable nutrients (Viitamäki et al., 2022). Similarly, *Bryobacter* (Kulichevskaya et al., 2010), *Acidothermus* (Viitamäki et al., 2022), and *Conexibacter* (Monciardini et al., 2003) synthesize polysaccharides and phytohormones that regulate biogeochemical cycles and promote plant growth (Kalam et al. 2020). In contrast, the genera *Flavobacterium* and *Sediminibacterium*, which demonstrated higher abundance under ambient conditions, play roles in carbon uptake and substrate biodegradation (Wilson et al., 2016). Their relative abundance in spring or summer seasons suggests their involvement in plant-microbe interactions that may influence carbon cycling in wild blueberry soils.

The cooperative interactions among these bacteria may contribute to the stability of microbial communities under warming conditions. For instance, *Acidothermus* exhibited positive associations with other taxa, such as *Xanthobacteraceae* and *Roseiarcus*, which are known to mediate biochemical processes that promote plant growth (Qu et al., 2020). As observed in previous studies, this microbial cooperation could also enhance plant resilience and biomass production under warming conditions (Cavaliere et al,. 2017).

## 5. Conclusions

This study represents an important step toward understanding the effects of global warming on the soil microbiome of wild blueberries. The results indicate that warming had a limited impact during the summer compared to the spring, likely due to the naturally elevated temperatures in the summer months. However, since the bacterial analyses in this study focused only on the spring and summer, it remains unclear how bacterial communities respond during the fall and winter when ambient temperatures are significantly lower. Despite this limitation, our findings show that warming influences bacterial communities in the early growing season of wild blueberry crops, suggesting a future shift in temporal dynamics in bacterial activity under global warming. Future research should explore the long-term effects of warming on soil bacterial and fungal composition and function across all four seasons in Maine, as well as the mechanistic linkage between bacterial seasonal activity and plant phenology, especially root dynamics.

## Acknowledgments

This project was funded by the USDA National Institute of Food and Agriculture (Hatch Project Number ME0-22021) through the Maine Agricultural and Forest Experiment Station, as well as by the University of Maine Faculty Summer Research Award, the Wild Blueberry Commission of Maine, Jasper Wyman & Son Inc., and the Maine Department of Agriculture, Conservation, and Forestry (SCBGP). We also thank Yu-Ying Chen, Marlys Rietdyk, and Nicholas Hershbine for their help with soil sample collection and DNA extraction and Jianjun Hao for technical assistance.

## Author contribution

OAA, SI, and YJZ conceived and designed the research. OAA conducted experiments, curated, analyzed and visualized data. OAA, SI, and YJZ interpreted data and supervised the project. OAA wrote the original manuscript. OAA, SI, LC, JH, YYC and YJZ revised and edited the manuscript. All authors read and approved the manuscript.

