## Supplemental Figure1_and_2 for "Warming treatments shift the temporal dynamics of diversity and composition of bacteria in wild blueberry soils"

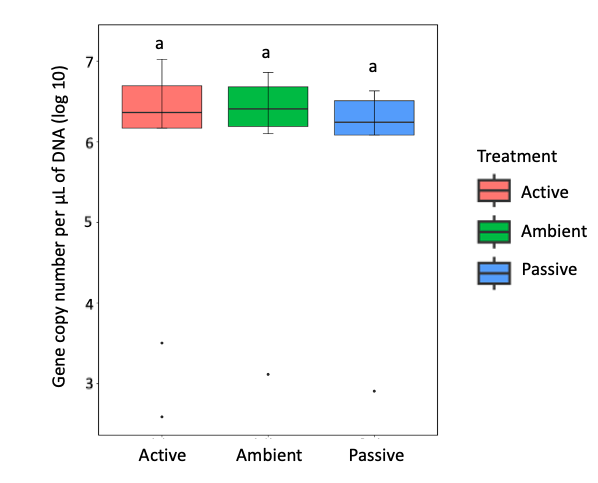

**Figure S1:** Absolute bacteria abundance determined by qPCR analysis of log 10-transformed 16S rRNA gene copies under the different treatments. Box topped with the same letters indicates no significant difference (*p* > 0.05).

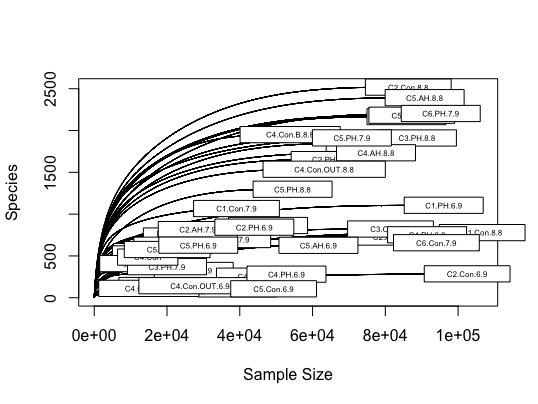

**Figure S2:** The rarefaction curves of bacterial sequence variants in wild blueberry soils under active warming (AH), passive warming (PH), and ambient temperatures (Con).
